# Transcriptional and mutational signatures of the aging germline

**DOI:** 10.1101/2021.11.22.469565

**Authors:** Evan Witt, Christopher B Langer, Nicolas Svetec, Li Zhao

**Affiliations:** Laboratory of Evolutionary Genetics and Genomics, The Rockefeller University, New York, NY 10065, USA

## Abstract

Aging is a complex biological process that is accompanied by changes in gene expression and mutational load. In many species, including humans, older fathers pass on more paternally-derived *de novo* mutations; however, the cellular basis and cell types driving this pattern are still unclear. To explore the root causes of this phenomenon, we performed single-cell RNA-sequencing (scRNA-seq) on testes from young and old male *Drosophila,* as well as genomic sequencing (DNA-seq) on somatic tissues from the same flies. We found that early germ cells from old and young flies enter spermatogenesis with similar mutational loads, but older flies are less able to remove mutations during spermatogenesis. Mutations in old cells may also increase during spermatogenesis. Our data reveal that old and young flies have distinct mutational biases. Many classes of genes show increased post-meiotic expression in the germlines of older flies. Late spermatogenesis-enriched genes have higher dN/dS than early spermatogenesis-enriched genes, supporting the hypothesis that late spermatogenesis is a source of evolutionary innovation. Surprisingly, young fly enriched genes show higher dN/dS than old fly enriched genes. Our results provide novel insights into the role of the germline in *de novo* mutation.

## Introduction

Aging is a process that is accompanied by phenotypic changes in animals. These phenotypic changes include both observable traits and intermediate traits, such as gene expression. Aging can also impact the health of offspring and evolution when it occurs in the reproductive period, such as passing a higher amount of *de novo* mutations to the offspring. Most novel mutations are inherited from the paternal germline, and the number of mutations inherited increases with paternal age (Crow, 2000; Gao et al., 2016, 2019). Some studies have attributed excess paternal mutations to the increased number of cell divisions that cycling spermatogonial stem cells undergo throughout the life of the male (Drost and Lee, 1995; Gao et al., 2011; Li et al., 1996). Conversely, other reports have found that the excess cell divisions do not track the ratio of maternal to paternal mutations during aging (Gao et al., 2019; Huttley et al., 2000), suggesting instead that lifestyle, chemical and environmental factors cause this discrepancy (Irigaray et al., 2007; Parkin et al., 2011). Previous studies of the effect of age on paternally inherited mutations have inferred *de novo* mutations through sequencing of parents and offspring (Gao et al., 2016). These methods are highly useful, but they only capture *de novo* mutations that have evaded repair mechanisms, ended up inside a viable gamete, fertilized an egg, and created a viable embryo. Much less is known about the dynamics of mutation and repair inside the male germline. One study found that mutations arise least frequently in human spermatogonia (Moore et al., 2021). In our previous work (Witt et al., 2019), however, we instead found that mutational load is highest in the earliest stages of spermatogenesis. Taken together, these results imply that most mutations occur prior to germline stem cell (GSC) differentiation and are removed during spermatogenesis. But are these mutations replicative in origin? If so, we would expect germline stem cells from older flies to be more mutated than those from younger flies.

In addition to these mutational effects, aging is known to cause other germline phenotypes such as lower numbers of germ cells and reduced germline stem cell proliferative capacity (Lee et al., 2020). The GSC microenvironment also undergoes chemical changes associated with reductions in fecundity (Jones, 2007). These phenotypic consequences, in fact, could be linked to mutation, as the germline mutational rate in young adults correlates with longevity (Cawthon et al., 2020). As such, the germline mutation rate has consequences for both an organism and its descendants.

In our previous study (Witt et al., 2019), we used single-cell RNA-sequencing (scRNA-seq) to follow germline mutations throughout *Drosophila* spermatogenesis and found evidence that germline mutations decline in abundance throughout spermatogenesis. We also found evidence that some germline genome maintenance genes are more highly expressed in GSCs and early spermatogonia, the earliest male germ cells. Our results were in line with the idea that active DNA repair plays a role in the male germline (Xia et al., 2020), but since age is an important factor for mutational load, in this study we directly compare patterns in young and old testis.

To study transcriptional and mutational signatures in the aging germline, we generated scRNA-seq data from *Drosophila melanogaster* testes 48 hours and 25 days after eclosion (“Young” and “Old” respectively). We also sequenced correlated genomic DNA from each sample to confirm that each detected mutation was a real *de novo* germline mutation. Our results support our previous observation that the proportion of mutated cells declines throughout spermatogenesis for young flies. For old flies, however, the proportion of mutated cells begins high and remains high throughout spermatogenesis. We found that on a molecular level, each class of older germ cell has a higher mutational burden than comparable cells from young flies. Additionally, older flies carry a higher proportion of C>G and C>A mutations. Our results indicate that the old germline is more highly mutated and transcriptionally dysregulated. We did not find evidence of a profound shift in the expression of genome maintenance genes; however, a number of these genes were highly expressed in young and old flies. We also find that patterns of global gene expression differ between young and old testes, including increased post-meiotic expression of *de novo* genes, TEs, and canonical genes. We found that early spermatogenesis-enriched genes have lower dN/dS than late spermatogenesis-enriched genes, and that genes enriched in older germ cells have lower dN/dS than genes enriched in younger germ cells. These findings provide a deeper insight into the process of spermatogenesis as a key source of *de novo* mutations.

## Results

### A cell atlas of the aged male germline

We aimed to capture representative cell types from the major somatic and germline cell types of the testis (Figure 1A). We generated testes scRNA-seq data from male flies 48 hours (young) and 25 days (old) after eclosion to facilitate the identification of *de novo* mutations (Figure 1B, Supplemental Table 1). Each of the six libraries was made with approximately 30 pairs of fly testes. We used Cellranger (Zheng et al., 2017) to align these libraries against the FlyBase (Thurmond et al., 2019) *D. melanogaster* genome (version R6.32). We used previously established marker genes (Witt et al., 2021a) to annotate cell types for the young and old flies separately. Dot plots showing the expression of key marker genes are shown in Supplemental Figure 1. To confirm that mutations observed are from the germline, we prepared and sequenced somatic genomic DNA libraries from the carcasses of the same flies used for scRNA-seq and used these samples as control.

**Figure 1:**
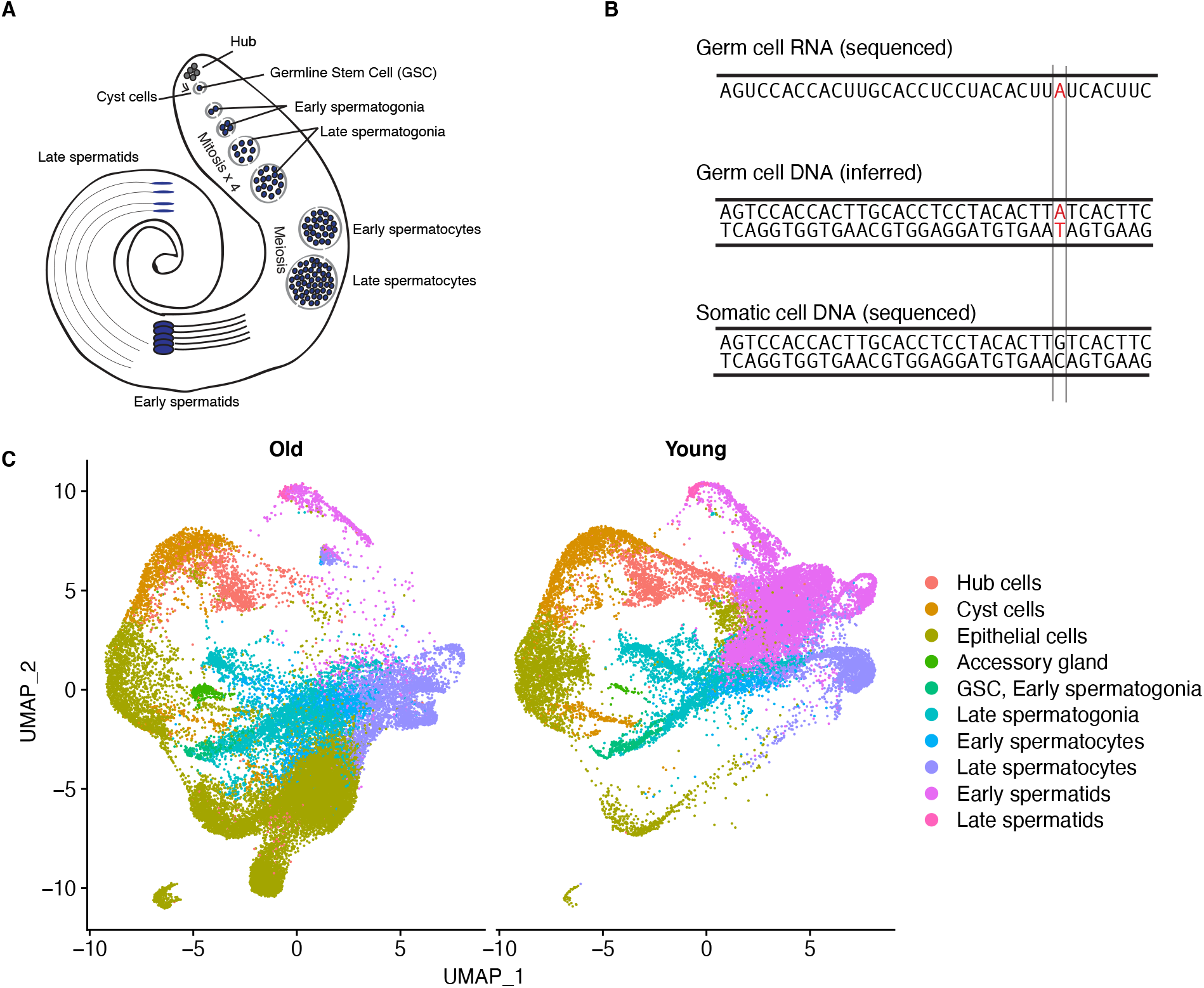
Overview of experimental design and visualization of old and young datasets. A) Diagram of *Drosophila* testis cell types and the marker genes used to identify each cell type. B) experimental rationale: we infer mutated genomic sites in germ cells using scRNA-seq data. If the same locus is unmutated in somatic cell DNA, we call the SNP in red a *de novo* mutation. We can detect a mutation if it is present on both strands or only the template strand. C) dimensional reduction showing the cell-type assignments of scRNA-seq data from young and old flies.

Using Seurat 4 (Satija et al., 2015), we classified somatic cells into four broad types: hub cells, cyst cells, accessory gland, and epithelial cells. We split germ cells into six types, listed from earliest to latest: germline stem cells/early spermatogonia, late spermatogonia, early spermatocytes, late spermatocytes, early spermatids, and late spermatids. In total, we characterized 23489 cells from young flies and 28861 cells from old flies (Figure 1C, Supplemental Table 1). We found that for each age group and cell type, the 3 replicates from each age group largely corroborate each other, with Pearson’s r values over 0.91 between replicates and cells of the same age (Supplemental Figure 2). After cell type assignments, we used Seurat 4 to perform downstream analyses on the integrated dataset.

### Older flies show impaired mutational repair during spermatogenesis

We identified germline SNPs in each sample and matched them to every cell with reads corroborating a given SNP (Figure 2). To assess the mutational burden between young and old flies, we compared, for each cell type, the proportion of cells where at least one mutation was detected. We also compared, for every cell type, the number of detected SNPs per unique molecular index (UMI) to account for differences in coverage between libraries. We did not find any SNPs in more than one replicate, indicating that recurrent age-related transcriptional errors or RNA editing events did not bias our results.

**Figure 2:**
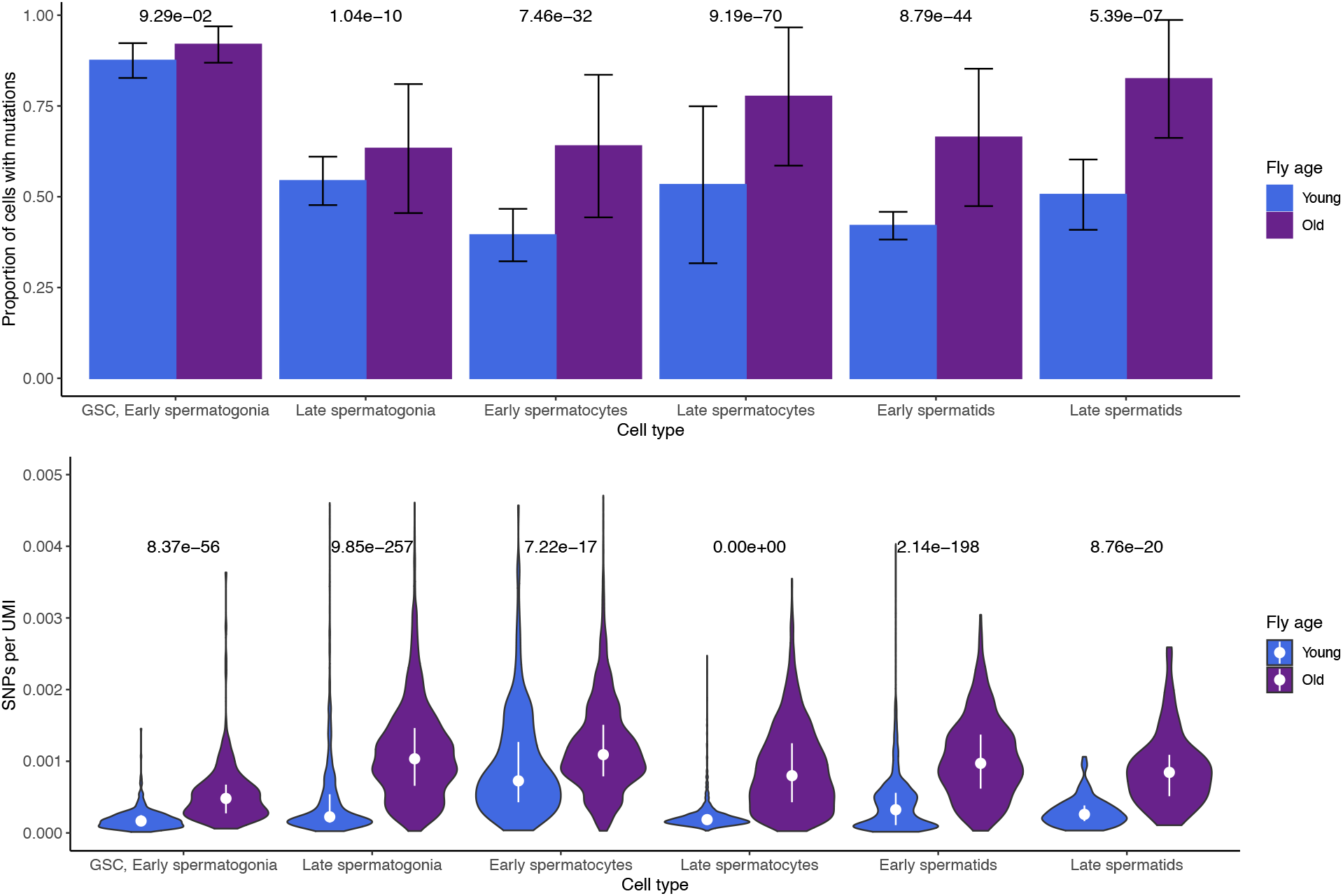
The proportion of mutated cells declines more for young flies than old flies, while old flies have a higher mutation load. A) For old and young flies and every cell type, shown are the proportions of cells of each type carrying at least one mutation. Error bars are standard error. P values are Bonferroni-corrected from a chisquare test of mean proportions between young and old cells of a type. The proportion of mutated cells declines more for young flies than not old flies, during spermatogenesis. B) for each cell, the number of SNPs divided by the number of Unique Molecular Indices (UMIs) detected, a proxy for read depth. P values are Bonferroni-corrected from a chi-square test of proportions comparing young and old cells of a type. Ever class of older germ cells has more mutations per RNA molecule detected. This indicates that the higher mutational load of older flies precedes spermatogenesis, and may increase during spermatogenesis.

In young flies, the proportion of mutated cells declines drastically during spermatogenesis, indicating that lesions are either repaired or that mutated cells are removed from the population. Old flies begin spermatogenesis with a similar proportion of mutated GSC/early spermatogonia, but their mutational burden remains high throughout spermatogenesis. Proportions of mutated cells are statistically similar for young and old flies in GSC/Early spermatogonia but begin to diverge in later cell types. By the end of spermatogenesis, young flies achieve a lower proportion of mutated cells compared to older flies, whose proportion of mutated cells remains high throughout.

Old spermatocytes and spermatids have significantly higher proportions of mutated cells (Figure 2A). To confirm that this trend was not confounded by different read depths across replicates, we counted the number of detected SNPs per Unique Molecular Index (UMI) for every cell type and found that RNA molecules from old cells are consistently more likely to carry mutations than young cells of the same type (Figure 2B). This suggests that much of the elevated mutational load of the older germline occurs before spermatogenesis, accumulated within cycling germ cells. However, we also observe that later germ cells from old flies have more mutations per RNA molecule than older GSC/early spermatogonia. This observation would be expected if some germline mutations arose during spermatogenesis.

### T>C and C>G substitutions are enriched in old flies

We compared the relative proportions of the six major classes of mutation between young and old flies. Young flies and old flies have distinct mutational signatures. In young flies, we found that C>T and T>G mutations are enriched compared to old flies (Figure 3). Using a chisquare test of proportions, we found that old flies were significantly enriched for T>C and C>G mutations. This suggests an age-related mutational or repair bias during spermatogenesis. We asked whether these mutational signatures were due to differential activity of genome maintenance genes (Svetec et al., 2016; Witt et al., 2019) and found that, as a group, genome maintenance genes are similarly expressed in most cell types (Supplemental Figure 3). As such, the mutational load of older flies cannot be purely attributed to age-related global downregulation of genomic maintenance genes, although it is likely that the regulation of genome maintenance genes at the protein level differs between young and old testis.

**Figure 3:**
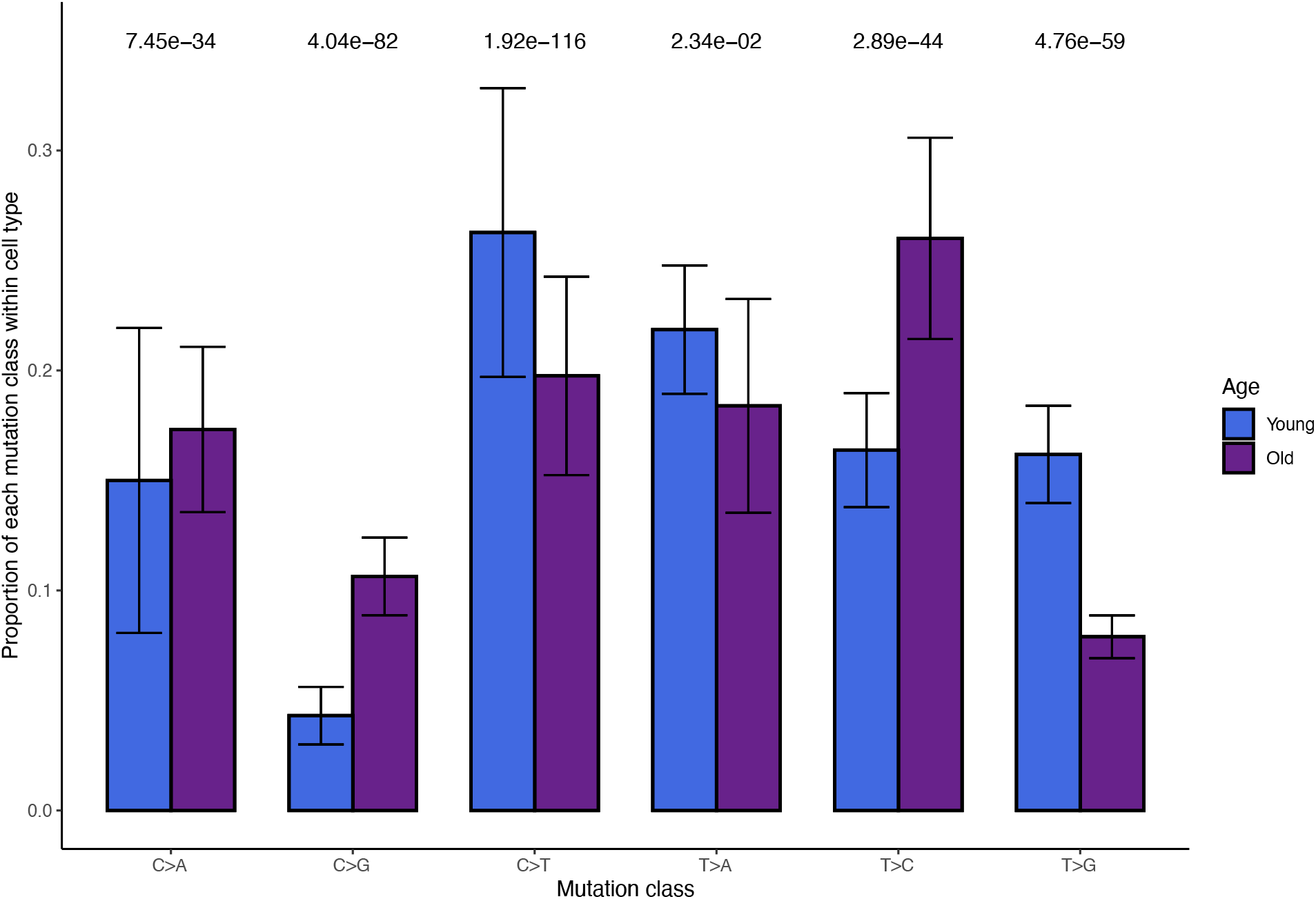
Age-related trends in mutational signatures. For young and old flies, shown are the relative proportions of the 6 types of mutations (each class is equivalent to a complementary mutation, for example, T>G also represents A>C). Error bars are standard error. Bonferroni-corrected P values are from a chi square test of proportions comparing the mean proportions between young and old flies. T>C and C>G mutations are enriched in old flies, while C>T and T>G mutations are underrepresented.

### Many genome maintenance genes are more highly expressed in early germ cells from young flies

We performed differential expression testing between every cell type (Table 1) and focused on a list of 211 genes related to DNA damage repair compiled from our previous work (Svetec et al., 2016; Witt et al., 2019). We found that in GSC/early spermatogonia, 7 genome maintenance genes were more highly expressed in young flies and 14 more highly expressed in old flies (Supplemental Figure 3, Supplemental Table 2). In spermatocytes and spermatids, comparatively few genes are differentially expressed between young and old testes. This corroborates our earlier observation that genome maintenance genes are generally less expressed in old GSC/early spermatogonia, which is also the most mutated cell type in both old and young flies. Depleted expression of genome maintenance genes in the earliest germ cells could impact the efficiency of germline DNA repair throughout spermatogenesis.

**Table 1:** Numbers of age-enriched genes per cell type. Arranged from top (early) to bottom (latest). For young flies, the cell type with the most enriched genes is GSC/Early spermatogonia, the earliest germ cell class. Late spermatids are the most mutated class of cell in older flies. In every class of cells except GSC, early and late spermatogonia, old flies have more enriched genes than young flies.

| Cell type | # genes enriched in young flies | # genes enriched in old flies |
| --- | --- | --- |
| **GSC, Early spermatogonia** | 273 | 251 |
| **Late spermatogonia** | 144 | 337 |
| **Early spermatocytes** | 40 | 367 |
| **Late spermatocytes** | 80 | 281 |
| **Early spermatids** | 167 | 277 |
| **Late spermatids** | 52 | 68 |

Two transcription-related genes, Rbp8 (FBgn0037121) and RpII15 (FBgn0004855) are more highly expressed in young GSC/Early spermatogonia than in old flies. Rbp8, also known as B52, is essential for DNA topoisomerase I recruitment to chromatin during transcription (Juge et al., 2010). DNApol-iota (FBgn0037554), which is a gene involved in translesion synthesis (Ishikawa et al., 2001) and may be important UV damage response (Svetec et al., 2016), was highly expresssed in young GSCs. This suggests that our observed mutational signatures of older flies might be caused by defects in transcription-coupled repair or DNA damage related repair, although the mechanisms of germline genomic surveillance are yet to be fully understoods. In old flies, lower expression of RpII15, also known as RNA Polymerase II, subunit I, might further explain reduced transcription in these cell types from old flies. The gene 14-3-3epsilon (FBgn0020238) is highly expressed in old GSC and spermatogonia, which may play a role in cell division and apoptosis in early germ cells for old testis (Su et al., 2001). The cellular-level mutational signature (Supplemental Figure 4) is in line with previous work suggesting that highly expressed genes evolve more slowly (Drummond et al., 2005; Good and Nachman, 2005).

### *De novo* genes, transposable elements, and other genes show increased post-meiotic expression in older flies

In our previous work we found that *de novo* genes are highly enriched in meiotic cells (Witt et al., 2019). We asked whether transcriptional dysregulation of the aging germline could impact the expression of genetic novelties such as *de novo* genes and transposable elements (TEs). We performed a parallel analysis of our scRNA-seq data with a custom reference containing 267 testis-expressed *de novo* genes identified from our 2019 study, as well as 239 TEs (Lawlor et al., 2021). We then scaled expression of every gene, centered at zero to compare the expression patterns of groups of genes (Figure 4).

**Figure 4:**
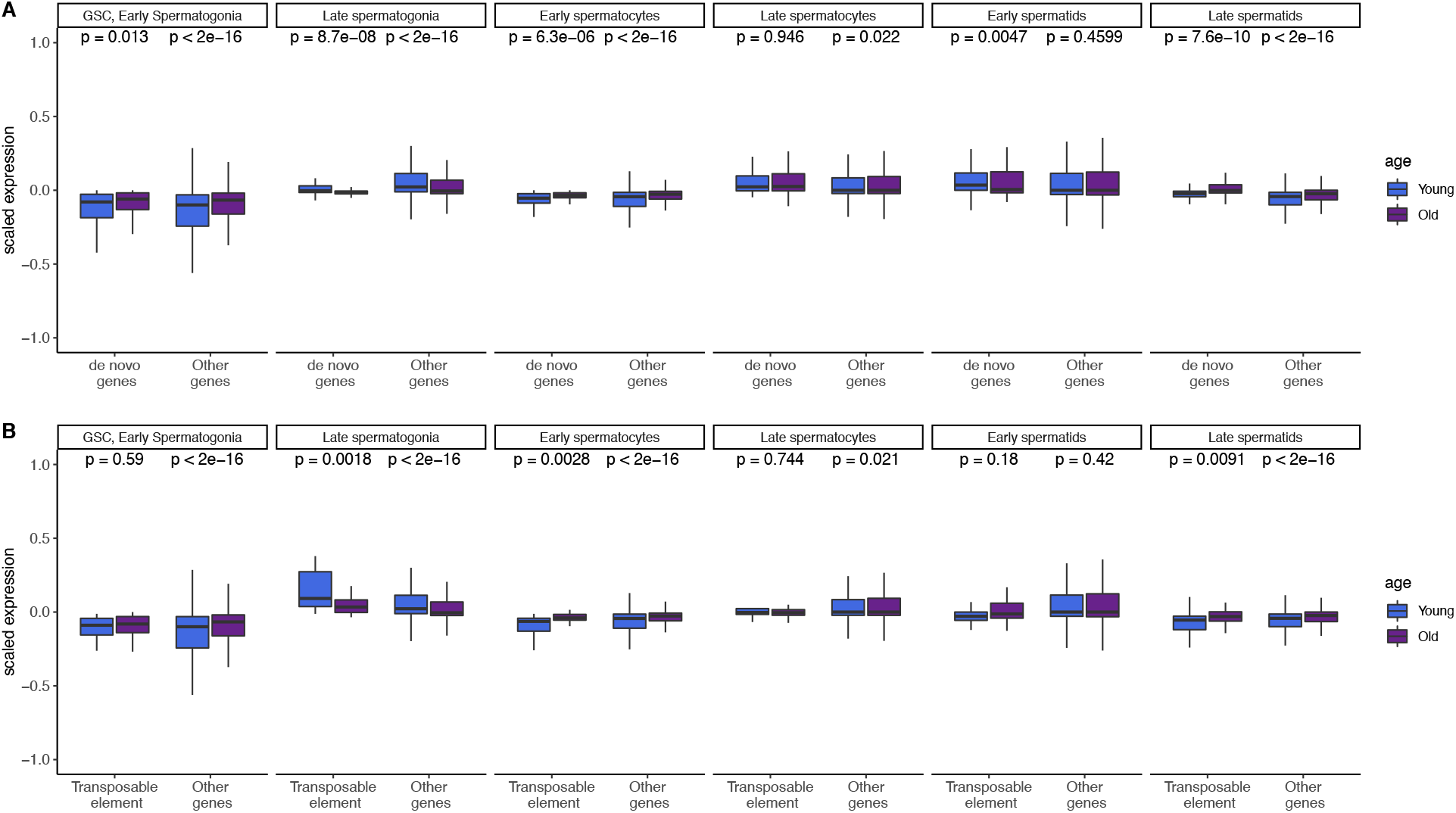
Global expression patterns of *de novo* genes and transposable elements changes with age in each cell type. A.) Scaled expression of *de novo* genes and other genes (not including TEs) across cell types. Expression of both gene types is enriched in the late spermatids of older flies. B.) Scaled expression of TEs and other genes (not including *de novo* genes) across cell types. Transposable element expression is highly enriched in late spermatogonia of young flies, and enriched in the late spermatids of older flies. Figure shows p values by Wilcoxon rank sum tests.

Overall, the expression patterns for *de novo* genes resemble those of other genes, and both *de novo* genes and other genes are more expressed in the late spermatids of older flies (Figure 4A, p = 7. 6e-10, p < 2e-16, Supplemental Table 3). This suggests that *de novo* genes show similar expression regulation related to aging compared to old genes and that the regulatory environment acts similarly to conserved and young genes.

We were also interested in the expression of TEs, since TE suppression is important for germ cell development (Lee and Langley, 2010). In other tissues, such as the aging brain, the amount of certain types of TEs change with age (Li et al., 2013). We found TE expression is also globally enriched in older late spermatids (Figure 4B, Supplemental Table 4), however this pattern is not more extreme compared to annotated genes. Since gene expression is supposed to stop after meiosis, it is possible that elevated gene expression after meiosis is a consequence of reduced post-meiotic transcriptional suppression. Although all 3 classes of genes were also more highly expressed in the late spermatogonia of young flies, this increase was strikingly large for transposable elements (p = 0.0018). This result corroborates earlier work which found a similar burst of transposon activity in early spermatogenesis, likely when spermatogonia transit to spermatocytes (Lawlor et al., 2021). As such, the reduced early TE expression in older germlines may reflect a dysregulated transcriptional environment.

### Early spermatogenesis-biased genes have lower dN/dS than late spermatogenesis-biased genes

We asked whether functional constraint varies for age-specific or cell-type specific genes. We defined “age-biased” genes as genes differentially expressed between old and young flies in the same cell type. We defined “stage-biased” genes as genes differentially expressed between cell types of the same age. To find stage-biased genes, we split our dataset into “old” and “young” cells and then performed Seurat’s FindMarkers function between GSC/early spermatogonia (early germline) and late spermatids (late germline) for each age group. Using dN/dS data from flyDIVaS (Stanley and Kulathinal, 2016), we compared dN/dS values for early-stage-biased and late-stage-biased genes. In both young and old flies, the genes with expression bias toward later stages exhibit higher dN/dS than genes biased in GSC/early spermatogonia (Figure 5A), which is in line with the idea that genes expressed in late spermatogenesis may evolve rapidly. Note that spermatocytes and spermatids are also hotspots for the expression of novel genes including *de novo* originated genes. Our findings are also similar to a murine study which found that genes expressed in early spermatogenesis are under more evolutionary constraint than genes expressed late germ cells (Schumacher and Herlyn, 2018).

**Figure 5:**
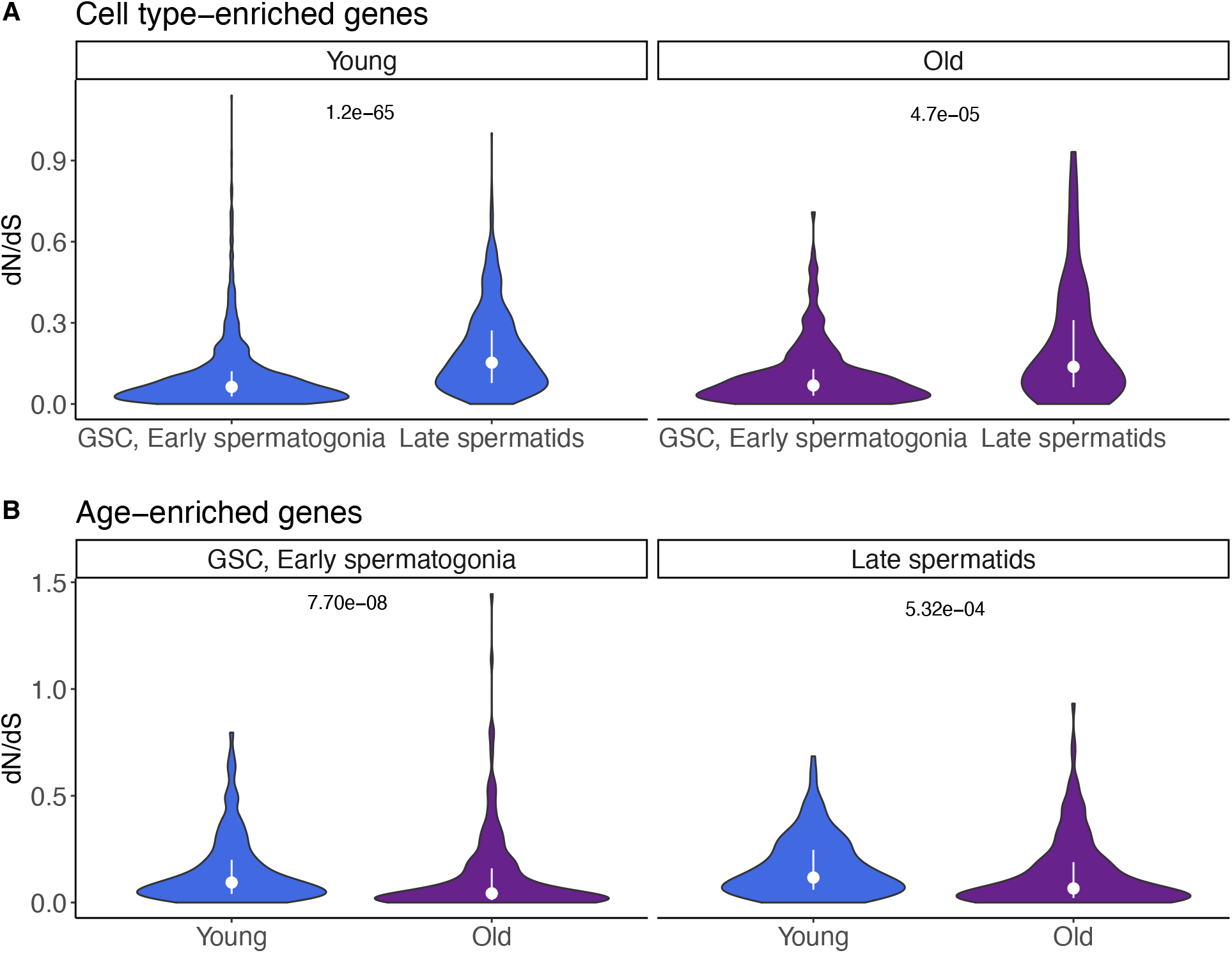
dN/dS trends of cell type-enriched (stage-biased) and age-enriched (age-biased) genes. We calculated gene enrichments in two different ways. First, we calculated gene enrichment between GSC/early spermatogonia and late spermatids separately for young and old flies. Then, we identified genes enriched in young and old cells within a cell type. A) Cell type-enriched genes: in both old and young flies, genes enriched in late spermatids have higher dN/dS than genes enriched in GSC/early spermatogonia (Wilcoxon rank sum test, p values 1.2e-65, 4.7e-5, respectively). B) Age-enriched genes: In GSC/early spermatogonia, but not late spermatids, young-enriched genes have higher dN/dS than old-enriched genes (Wilcoxon rank sum test, p values 7.70e-08 and 5.32e-04, respectively). dN/dS values are from flyDIVaS.

To identify age-biased genes, we divided the dataset by cell type and identified genes enriched in old or young cells of the same type. We found that in both GSC/early spermatogonia and late spermatids, genes biased in young cells have higher dN/dS than genes biased in old cells (Figure 5B). This result show that many early-germ-cell-biased genes in young flies evolve rapidly, that is, a higher proportion of rapidly evolving genes are enriched in young-age germline stem cells and spermatogonia than in old ones. This suggests that, if antagonistic pleiotropy in aging plays an important role in the shift of gene expression (Austad and Hoffman, 2018; Williams, 1957), rapidly evolving genes are often expressed and function in young animals and are subject to antagonistic pleiotropy.

## Discussion

Mutational load is an equilibrium between mutation and repair. In old flies, this equilibrium may shift away from repair. In this work, we show that the germline of older flies is less able to remove *de novo* mutations compared to the germline of young flies. Throughout spermatogenesis, we observed that germ cells from older flies have more mutations per RNA molecule than comparable cells from younger flies. This finding adds a new explanation for the still-controversial mechanism behind the increased age-dependent mutational load of the male germline. Our finding of increased mutational load in older GSC/early spermatogonia suggests that much of the mutational load of older flies accumulates prior to spermatogenesis. However, later germ cells of older flies have more SNPs per UMI, suggesting that mutations may accumulate during spermatogenesis. Our work corroborates previous work that found that the huge excess of male germline divisions is too large to explain the much smaller ratio of male/female-inherited mutations during parental aging (Gao et al., 2019). Our finding that early germ cells from young and old flies are similarly mutated supports the notion that many age-related germline mutations are not due to replicative processes. Some of our conclusions are in line with recent findings that germline stem cells have the lowest mutation rate of any human cell type (Moore et al., 2021).

In addition to being highly mutated, the older germline shows distinct mutational signatures compared to the younger germline. For example, we found a statistical overrepresentation of C>A and C>G mutations in old flies and an underrepresentation of C>T mutations. These altered ratios of single nucleotide polymorphisms could be caused by differential activity of DNA repair pathways in the old germline. We did not find strong evidence of global downregulation of genome maintenance gene expression but found lower expression of a few key transcriptional genes in GSC of older flies. Altered types and numbers of *de novo* mutations would likely have implications for a population, affecting the type and frequency of genetic novelties that emerge (Loewe and Hill, 2010). In the future, it would be interesting to understand the molecular mechanisms contributing to mutational bias in germ cells. Additionally, these methods should be reproduced with mated flies to examine if germline mutational bias can be affected by a male’s reproductive activity.

We observed that scaled expression of all genes is generally down in GSC/early spermatogonia but up in late spermatids. The latter result is intriguing because transcription largely ceases after meiosis in the male *Drosophila* germline (Barreau et al., 2008). While the downregulation of transcription in early germ cells could have implications for germline DNA repair, the potential effects of increased post-meiotic transcription are less clear. It could have no effect, or it could affect spermatid maturation or sperm competition, potentially affecting fertility. Indeed, increased male age associates with reduced fertility in humans (Harris et al., 2011).

Other studies have proposed that the testis uses ubiquitous gene expression to detect genomic lesions and repair them with transcription-coupled repair (Xia and Yanai, 2022; Xia et al., 2020). Due to our bias towards detecting mutations in expressed genes, our data is not ideal to test this hypothesis. We noted, however, that genes with many detectable SNPs tend to be lowly expressed across replicates (Supplemental Figure 4), a finding that appears to be consistent with the transcriptional scanning model. Broadly, this is in line with previous work that highly expressed genes show lower dN/dS, which previous work show that this was driven by various types of selective pressures (Drummond et al., 2005; Good and Nachman, 2005). Our results surprisingly found this pattern in mutational bias, even before natural selection acts on the genes. This result also supports the notion that it is very unlikely that the germline mutations we observed are directly from transcription errors, as that would lead to more mutations to highly expressed genes. Impaired mutational repair of the older male germline may be partially due to deficiencies in transcription-coupled repair (Deger et al., 2019).

The global post-meiotic upregulation of transcription extends beyond conserved genes. We observed that many transposable elements are highly enriched in the early germline of young flies. Not only could this have implications for transposable element mobilization, it could also create heritable changes in chromatin structure, signaling, or gene expression (Chuong et al., 2017; Lanciano and Cristofari, 2020).

The global deregulation of gene expression during aging also has interesting implications for evolution. Consistent with prior work (Schumacher and Herlyn, 2018), we found that genes enriched in late germ cells have higher dN/dS than genes enriched in early germ cells for both young and old flies. This result suggests that spermatocytes and spermatids are sources of rapid evolution or positive selection. Considering that spermatocytes and spermatids are also the stages where *de novo* gene are the most abundant and where they likely function (Witt et al., 2019) our results highlight the importance of late spermatogenesis in transcriptional and functional innovation.

Unexpectedly, we also found that, within analogous cell types, genes enriched in cells from young flies have higher dN/dS than genes enriched in old flies. There are two possible explanations. First, rapidly evolving genes or genes with more adaptive changes may provide a greater evolutionary advantage to young flies than old flies. Second, since we used the scaled expression, old-enriched genes may reflect a specific set of genes that are essential for aging animals, which may be more likely to be house-keeping genes. Our results are interesting in the light of recent work which found that genes expressed later in life tend to fix nonsynonymous mutations more frequently (Cheng and Kirkpatrick, 2021). One should note that their methodology is different: their age-biased genes were identified from whole-body data, whereas ours were calculated just from male germ cells. Gene expression in the testis is often an outlier compared to other tissues (Witt et al., 2021b), so the results of these two studies are not necessarily in conflict. Nevertheless, the consistent pattern between this study and that of Cheng and Kirkpatrick (Cheng and Kirkpatrick, 2021) is that that genes enriched in late spermatids have a higher dN/dS than those enriched in early germ cells. In this way, at both whole-organism development and germline development level, late-stage biased genes tend to evolve more rapidly.

Our study design limits the detection of mutations in expressed transcripts. While we have strict criteria for the identification of novel SNPs, the abundance of false negatives could vary between cell types due to cell-specific variation in transcriptional activity. We are reassured because the most commonly mutated cell type in our datasets is GSC/early spermatogonia, consistent with our previous observations using different datasets and slightly different analytical pipelines. If transcriptional activity biased our inference of mutational load, we would expect spermatocytes, the most transcriptionally active cell type, to appear the most mutated instead. This potential confounder would be resolved if a method became available to simultaneously perform RNA-seq and whole-genome sequencing on thousands of single cells. Current methods may not capture enough cells to comprehensively profile rarer cell types, but we expect this technical challenge to improve with future technological advances.

Another potential confounder is that aging might create the appearance of germline SNPs through reduced transcriptional fidelity (Verheijen and van Leeuwen, 2017). We do not think this is a significant source of error, since our SNPs are verified by multiple independent reads and are not allowed to be present in more than one dataset, in fact, we did not observe mutations occur in multiple datasets after filtering, suggesting that hidden common transcription errors or RNA editing do not impact our work. To understand the role that age-related transcriptional fidelity significantly plays in our results, this topic would benefit from high throughput combined scRNA/DNA sequencing from the same cells. The technology allowing us to trace *de novo* mutations throughout the germline is still very new, and we look forward to technological advancements in this exciting field.

## Methods

### ScRNA-seq library preparation and sequencing

In this study, we used young and old RAL517 for experiments. Briefly, flies were reared in the standard corn syrup medium at room temperature with synchronized 12:12 hour light:dark cycle. Both age groups of flies were kept as virgins after eclosion. Young and old samples were collected 48 hours and 25 days after eclosion, respectively. In detail, virgin males were collected within one hour after eclosion and were transferred to new vials. After 48 hours, we dissected 30 pairs of young testes for single-cell suspension and scRNA-seq and kept the carcasses for DNA sequencing. After 25 days, we dissected 30 pairs of old testes for single-cell suspension and scRNA-seq and kept the carcasses for DNA sequencing. We generated three biological replicates for young and old flies, respectively. Flies were all dissected in the morning (ZT1-ZT3 in our lab environment) to reduce expression fluctuation due to circadian rhythm. Single-cell testis suspensions were prepared as described in our previous work (Witt et al., 2019). Libraries were prepared with 10X Chromium 3’ V3 kit and sequenced with Illumina Hiseq 4000.

### Genomic DNA preparation

Fly carcasses were frozen at −80°C, then ground in 200 μL 100 mM Tris-HCl, pH 7.5, 100 mM EDTA, 100mM NACl, 0.5% SDS. The mixtures were incubated at 65°C for 40 minutes. Then, 160 μl KAc and 240 μl 6M LiCl were added, tubes were inverted 10 times and placed on ice for 30 minutes. Samples were then centrifuged at 18000g at 4°C for 15 minutes. The supernatant was transferred into a new tube and an equivalent volume of isopropanol was added and mixed by inversion. Samples were spun for 15 minutes at 18000g, and the supernatant was discarded. Pellets were washed with 800 μL 70% ethanol and samples were spun at 18000g for 5 minutes, and supernatant discarded. Pellet was air dried for 5 minutes and resuspended in 100 μL nuclease-free water. DNA was then sent for Illumina library preparation and sequencing by Novogene.

### ScRNA-seq data processing

ScRNA-seq data were aligned with Cellranger Count and further processed with Seurat. To assign cell types with Seurat, we used marker genes described in Witt et al. 2021 (Witt et al., 2021a). We normalized the replicates with SCTransform in Seurat and integrated them into a combined Seurat object. We performed clustering and annotation together on all replicates, using normalized counts from the “SC T” slot. Cells were annotated based on the expression patterns of marker genes. Clusters enriched in *bam* and *aub* are GSC/early spermatogonia(Kawase, 2004; Witt et al., 2019, 2021a), and adjacent clusters with less *bam/aub* and less *His2Av* are late spermatogonia. Clusters enriched in *fzo* and *twe* are early and late spermatocytes, respectively (Courtot et al., 1992; Hwa et al., 2002). Clusters with enriched *soti,* but not p-cup are early spermatids (Barreau et al., 2008) and clusters enriched in *p-cup* are late spermatids (Barreau et al., 2008). Clusters enriched in *MtnA* are somatic cells, *dlg1* defines cyst cells (Papagiannouli and Mechler, 2009) and *Fas3* defines hub cells. Epithelial cells are enriched in *MtnA* but not *Rab11* or *Fas3* (Witt et al., 2021a).

SNPs were called with bcftools (Narasimhan et al., 2016) separately for each single-cell library and each gDNA library. Per-base coverage was calculated for every gDNA sample with Samtools (Li et al., 2009). For each young and old scRNA-seq library, bcftools isec was used to extract mutations only present in the SC data and not the somatic gDNA. Using Samtools, we identified every cell barcode in the scRNA-seq data that corroborated every SNP (details in accompanying code). For each mutated position, we then verified that the corresponding locus in the gDNA file had at least 10 reads supporting the reference allele, and 0 reads supporting the alternative allele. We also required that every SNP be present only in a single scRNA-seq dataset, to reduce the chance that RNA editing events or transcription errors caused us to infer a SNP incorrectly. We also required every SNP to have >=2 reads corroborating it, reducing the potential impact of sequencing errors.

### Comparisons using scaled expression

To compare gene expression for groups of genes across replicates, we scaled expression using the ScaleData Seurat function separately on each replicate. Expression is scaled such that 0 represents a gene’s median expression across all cells, 1 represents 1 standard deviation above that gene’s mean expression, and −1 represents 1 standard deviation below. Within a cell type, each gene’s scaled expression was averaged between cells. Groups of genes were compared using a two-sample Wilcoxon rank sum test, and p values were adjusted with Bonferroni’s correction.

### Differential expression testing

For each germ cell type, we made a subset Seurat object containing just that cell type with old and young flies, assigning “age” as the cell identifier. We then used Seurat’s FindMarkers function with ident.1 as “Young” and ident.2 as “Old”. We classified genes with a Bonferroni-adjusted p value < 0.05 and Log2 fold change > 0 as enriched in young, and <0 as enriched in old. We then constructed volcano plots with the EnhancedVolcano package, while including differentially expressed genes from our list of 211 genome maintenance genes from our previous paper (Witt et al., 2019).

### *De novo* gene and TE analysis

*De novo* genes from our previous paper (Witt et al., 2019) were added to a reference GTF containing transposable elements from another study (Lawlor et al., 2021). This alternate reference was used to align reads from all libraries with Cellranger. Cell-type annotations were copied from the annotations made for the main Seurat object. Enriched *de novo* genes and TEs were detected with the FindMarkers function in Seurat.

### Data availability

Code used for processing of data is deposited at https://github.com/LiZhaoLab/Mutation_project. This repository will include permanent links to large data files including a Seurat RDS and mutation database. Raw sequence data will be released on SRA upon publication.

## Acknowledgements

We thank Hong Duan and Connie Zhao at Genomics Resource Center of Rockefeller University for their help with the scRNA-seq libraries, and members of Zhao lab for their helpful comments and suggestions.

## Reference

Austad, S.N., and Hoffman, J.M. (2018). Is antagonistic pleiotropy ubiquitous in aging biology? Evol. Med. Public Heal. 2018, 287–294.

Barreau, C., Benson, E., Gudmannsdottir, E., Newton, F., and White-Cooper, H. (2008). Post-meiotic transcription in *Drosophila* testes. Development 135, 1897–1902.

Cawthon, R.M., Meeks, H.D., Sasani, T.A., Smith, K.R., Kerber, R.A., O’Brien, E., Baird, L., Dixon, M.M., Peiffer, A.P., Leppert, M.F., et al. (2020). Germline mutation rates in young adults predict longevity and reproductive lifespan. Sci. Rep. 10, 10001.

Cheng, C., and Kirkpatrick, M. (2021). Molecular evolution and the decline of purifying selection with age. Nat. Commun. 12, 2657.

Chuong, E.B., Elde, N.C., and Feschotte, C. (2017). Regulatory activities of transposable elements: from conflicts to benefits. Nat. Rev. Genet. 18, 71–86.

Courtot, C., Fankhauser, C., Simanis, V., and Lehner, C.F. (1992). The *Drosophila cdc25* homolog *twine* is required for meiosis. Development 116, 405–416.

Crow, J.F. (2000). The origins, patterns and implications of human spontaneous mutation. Nat. Rev. Genet. 1, 40–47.

Deger, N., Yang, Y., Lindsey-Boltz, L.A., Sancar, A., and Selby, C.P. (2019). Drosophila, which lacks canonical transcription-coupled repair proteins, performs transcription-coupled repair. J. Biol. Chem. 294, 18092–18098.

Drost, J.B., and Lee, W.R. (1995). Biological basis of germline mutation: Comparisons of spontaneous germline mutation rates among drosophila, mouse, and human. Environ. Mol. Mutagen. 25, 48–64.

Drummond, D.A., Bloom, J.D., Adami, C., Wilke, C.O., and Arnold, F.H. (2005). Why highly expressed proteins evolve slowly. Proc. Natl. Acad. Sci. U. S. A. 102, 14338–14343.

Gao, J.-J., Pan, X.-R., Hu, J., Ma, L., Wu, J.-M., Shao, Y.-L., Barton, S.A., Woodruff, R.C., Zhang, Y.-P., and Fu, Y.-X. (2011). Highly variable recessive lethal or nearly lethal mutation rates during germ-line development of male Drosophila melanogaster. Proc. Natl. Acad. Sci. U. S. A. 108, 15914–15919.

Gao, Z., Wyman, M.J., Sella, G., and Przeworski, M. (2016). Interpreting the dependence of mutation rates on age and time. PLoS Biol. 14, 1–16.

Gao, Z., Moorjani, P., Sasani, T.A., Pedersen, B.S., Quinlan, A.R., Jorde, L.B., Amster, G., and Przeworski, M. (2019). Overlooked roles of DNA damage and maternal age in generating human germline mutations. Proc. Natl. Acad. Sci. 116, 9491 LP–9500.

Good, J.M., and Nachman, M.W. (2005). Rates of protein evolution are positively correlated with developmental timing of expression during mouse spermatogenesis. Mol. Biol. Evol. 22, 1044–1052.

Harris, I.D., Fronczak, C., Roth, L., and Meacham, R.B. (2011). Fertility and the aging male. Rev. Urol. 13, e184–e190.

Huttley, G.A., Jakobsen, I.B., Wilson, S.R., and Easteal, S. (2000). How important is DNA replication for mutagenesis? Mol. Biol. Evol. 17, 929–937.

Hwa, J.J., Hiller, M.A., Fuller, M.T., and Santel, A. (2002). Differential expression of the Drosophila mitofusin genes fuzzy onions (fzo) and dmfn. Mech. Dev. 116, 213–216.

Irigaray, P., Newby, J.A., Clapp, R., Hardell, L., Howard, V., Montagnier, L., Epstein, S., and Belpomme, D. (2007). Lifestyle-related factors and environmental agents causing cancer: An overview. Biomed. Pharmacother. 61, 640–658.

Ishikawa, T., Uematsu, N., Mizukoshi, T., Iwai, S., Iwasaki, H., Masutani, C., Hanaoka, F., Ueda, R., Ohmori, H., and Todo, T. (2001). Mutagenic and Nonmutagenic Bypass of DNA Lesions byDrosophila DNA Polymerases dpolη and dpolι. J. Biol. Chem. 276, 15155–15163.

Jones, D.L. (2007). Aging and the germ line: where mortality and immortality meet. Stem Cell Rev. 3, 192–200.

Juge, F., Fernando, C., Fic, W., and Tazi, J. (2010). The SR Protein B52/SRp55 Is Required for DNA Topoisomerase I Recruitment to Chromatin, mRNA Release and Transcription Shutdown. PLOS Genet. 6, e1001124.

Kawase, E. (2004). Gbb/Bmp signaling is essential for maintaining germline stem cells and for repressing bam transcription in the *Drosophila* testis. Development 131, 1365–1375.

Lanciano, S., and Cristofari, G. (2020). Measuring and interpreting transposable element expression. Nat. Rev. Genet. 21, 721–736.

Lawlor, M.A., Cao, W., and Ellison, C.E. (2021). A transposon expression burst accompanies the activation of Y-chromosome fertility genes during Drosophila spermatogenesis. Nat. Commun. 12, 6854.

Lee, Y.C.G., and Langley, C.H. (2010). Transposable elements in natural populations of Drosophila melanogaster. Philos. Trans. R. Soc. B Biol. Sci. 365, 1219–1228.

Lee, M.-H., Luo, H.-R., Bae, S.H., and San-Miguel, A. (2020). Genetic and Chemical Effects on Somatic and Germline Aging. Oxid. Med. Cell. Longev. 2020, 4684890.

Li, H., Handsaker, B., Wysoker, A., Fennell, T., Ruan, J., Homer, N., Marth, G., Abecasis, G., Durbin, R., and Subgroup, 1000 Genome Project Data Processing (2009). The Sequence Alignment/Map format and SAMtools. Bioinformatics 25, 2078–2079.

Li, W., Prazak, L., Chatterjee, N., Grüninger, S., Krug, L., Theodorou, D., and Dubnau, J. (2013). Activation of transposable elements during aging and neuronal decline in Drosophila. Nat. Neurosci. 16, 529–531.

Li, W.H., Ellsworth, D.L., Krushkal, J., Chang, B.H., and Hewett-Emmett, D. (1996). Rates of nucleotide substitution in primates and rodents and the generation-time effect hypothesis. Mol. Phylogenet. Evol. 5, 182–187.

Loewe, L., and Hill, W.G. (2010). The population genetics of mutations: good, bad and indifferent. Philos. Trans. R. Soc. B Biol. Sci. 365, 1153–1167.

Moore, L., Cagan, A., Coorens, T.H.H., Neville, M.D.C., Sanghvi, R., Sanders, M.A., Oliver, T.R.W., Leongamornlert, D., Ellis, P., Noorani, A., et al. (2021). The mutational landscape of human somatic and germline cells. Nature 597, 381–386.

Narasimhan, V., Danecek, P., Scally, A., Xue, Y., Tyler-Smith, C., and Durbin, R. (2016). BCFtools/RoH: a hidden Markov model approach for detecting autozygosity from next-generation sequencing data. Bioinformatics 32, 1749–1751.

Papagiannouli, F., and Mechler, B.M. (2009). discs large regulates somatic cyst cell survival and expansion in Drosophila testis. Cell Res. 19, 1139–1149.

Parkin, D.M., Boyd, L., and Walker, L.C. (2011). 16. The fraction of cancer attributable to lifestyle and environmental factors in the UK in 2010. Br. J. Cancer 105, S77–S81.

Satija, R., Farrell, J.A., Gennert, D., Schier, A.F., and Regev, A. (2015). Spatial reconstruction of single-cell gene expression data. Nat. Biotechnol. 33, 495–502.

Schumacher, J., and Herlyn, H. (2018). Correlates of evolutionary rates in the murine sperm proteome. BMC Evol. Biol. 18, 35.

Stanley, C.E.J., and Kulathinal, R.J. (2016). flyDIVaS: A Comparative Genomics Resource for Drosophila Divergence and Selection. G3 (Bethesda). 6, 2355–2363.

Su, T.T., Parry, D.H., Donahoe, B., Chien, C.T., O’Farrell, P.H., and Purdy, A. (2001). Cell cycle roles for two 14-3-3 proteins during Drosophila development. J. Cell Sci. 114, 3445–3454.

Svetec, N., Cridland, J.M., Zhao, L., and Begun, D.J. (2016). The Adaptive Significance of Natural Genetic Variation in the DNA Damage Response of Drosophila melanogaster. PLoS Genet. 12, e1005869.

Thurmond, J., Goodman, J.L., Strelets, V.B., Attrill, H., Gramates, L.S., Marygold, S.J., Matthews, B.B., Millburn, G., Antonazzo, G., Trovisco, V., et al. (2019). FlyBase 2.0: the next generation. Nucleic Acids Res. 47, D759–D765.

Verheijen, B.M., and van Leeuwen, F.W. (2017). Commentary: The landscape of transcription errors in eukaryotic cells. Front. Genet. 8, 219.

Williams, G.C. (1957). Pleiotropy, Natural Selection, and the Evolution of Senescence. Evolution (N. Y). 11, 398.

Witt, E., Benjamin, S., Svetec, N., and Zhao, L. (2019). Testis single-cell RNA-seq reveals the dynamics of de novo gene transcription and germline mutational bias in Drosophila. Elife 8, e47138.

Witt, E., Shao, Z., Hu, C., Krause, H.M., and Zhao, L. (2021a). Single-cell RNA-sequencing reveals pre-meiotic X-chromosome dosage compensation in Drosophila testis. PLOS Genet. 17, e1009728.

Witt, E., Svetec, N., Benjamin, S., and Zhao, L. (2021b). Transcription Factors Drive Opposite Relationships between Gene Age and Tissue Specificity in Male and Female Drosophila Gonads. Mol. Biol. Evol. 38, 2104–2115.

Xia, B., and Yanai, I. (2022). Gene expression levels modulate germline mutation rates through the compound effects of transcription-coupled repair and damage. Hum. Genet. 141, 1211–1222.

Xia, B., Baron, M., Yan, Y., Wagner, F., Kim, S.Y., Keefe, D.L., Alukal, J.P., Boeke, J.D., and Yanai, I. (2020). Widespread transcriptional scanning in testes modulates gene evolution rates. Cell 248–262.

Zheng, G.X.Y., Terry, J.M., Belgrader, P., Ryvkin, P., Bent, Z.W., Wilson, R., Ziraldo, S.B., Wheeler, T.D., McDermott, G.P., Zhu, J., et al. (2017). Massively parallel digital transcriptional profiling of single cells. Nat. Commun. 8, 14049.

