## Supplementary figures and tables for "Transcriptional and mutational signatures of the aging germline"

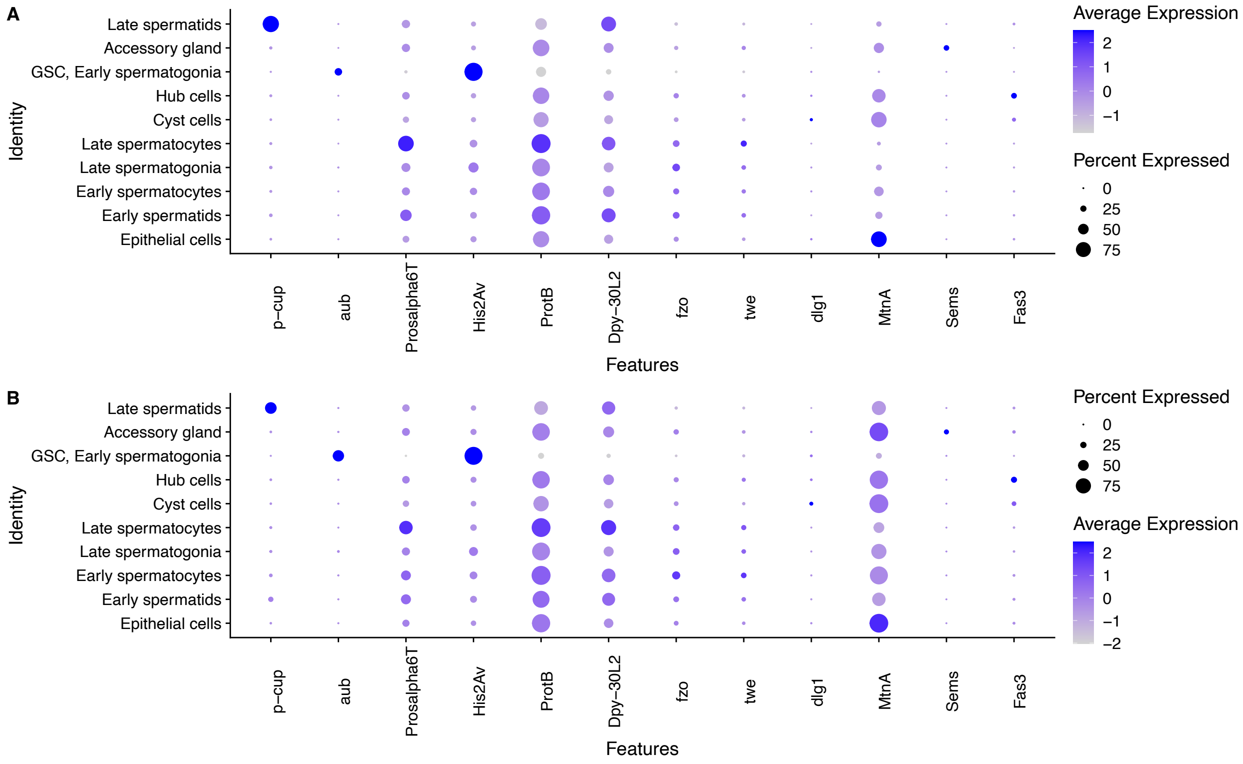


**Supplemental Figure 1: Dot plots of key marker genes in old, young fly testes.**

Split by cell type, these are the average expression values of the "SCT" slot in the old (A) and young (B) Seurat objects. Color corresponds to the level of expression, and the size of the dot represents the percent of cells of a class where a gene is detected. For example, *p-cup* was used to assign cells as late spermatids. *Bam* (Ohlstein and McKearin, 1997; Witt et al., 2019)*, aub* (Rojas-Ríos et al., 2017) and *vas* (Liu et al., 2009) were used to assign cells as GSC/early spermatogonia. *MtnA* (Faisal et al., 2014) was used to assign cells as epithelial cells, and *dlg1* (Papagiannouli and Mechler, 2009) was used to assign cyst cells. *Dpy-30L2* (Vardanyan et al., 2008) was used to differentiate early spermatids from late spermatocytes, and *fzo* (Hwa et al., 2002) was used to assign early spermatocytes. Late spermatogonia were assigned with *aub, bam, vas*, while having less *His2Av* (Jayaramaiah Raja and Renkawitz-Pohl, 2005) than GSC/early spermatogonia. Cells enriched with Sems were determined to be accessory gland.

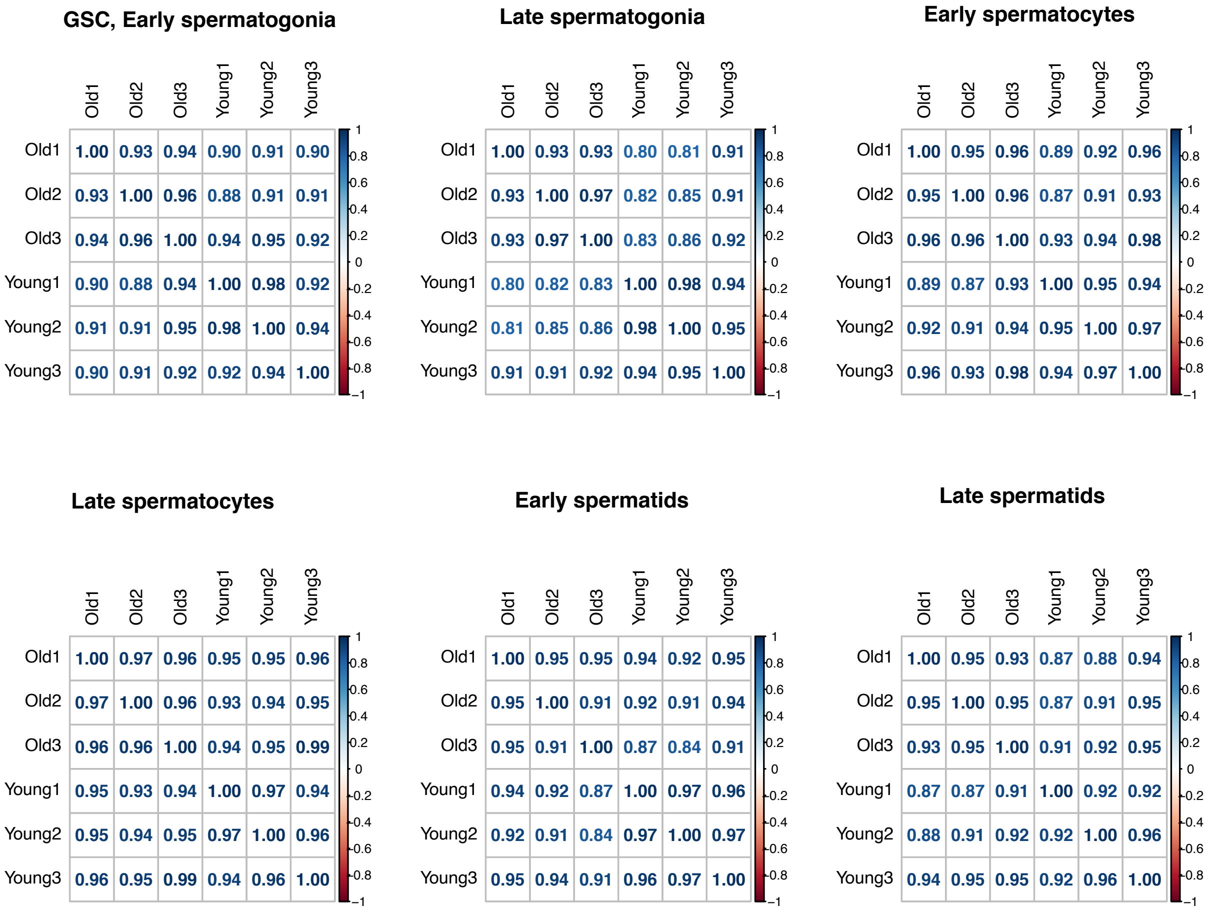


**Supplemental Figure 2: Correlograms of germ cells between scRNA-seq replicates.**

Correlations have been split between cell types. For each cell type, replicates from each age group all correlate with Pearson's R >0.91. Correlations were drawn from gene expression values from the "RNA" slot of the Seurat object using the corrplot R package.

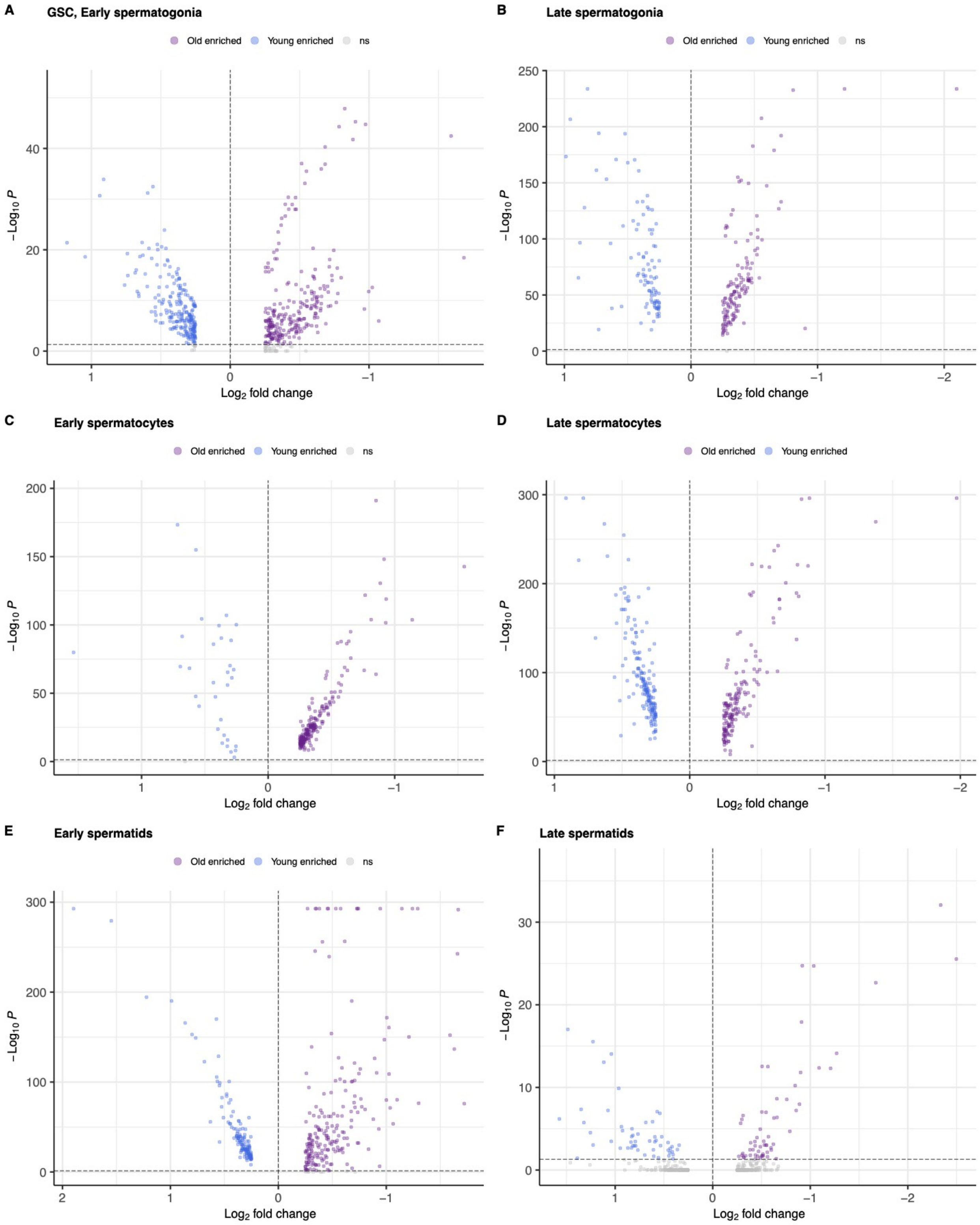


**Supplemental Figure 3: Age-related differential expression of genes, including genome maintenance genes.**

Shown are the results of differential expression tests between old and young flies, calculated separately for each cell type. Log2 fold changes refer to the ratio between expression in young compared to old flies. Enrichment statistics for genome maintenance genes are in Supplemental Table 2.

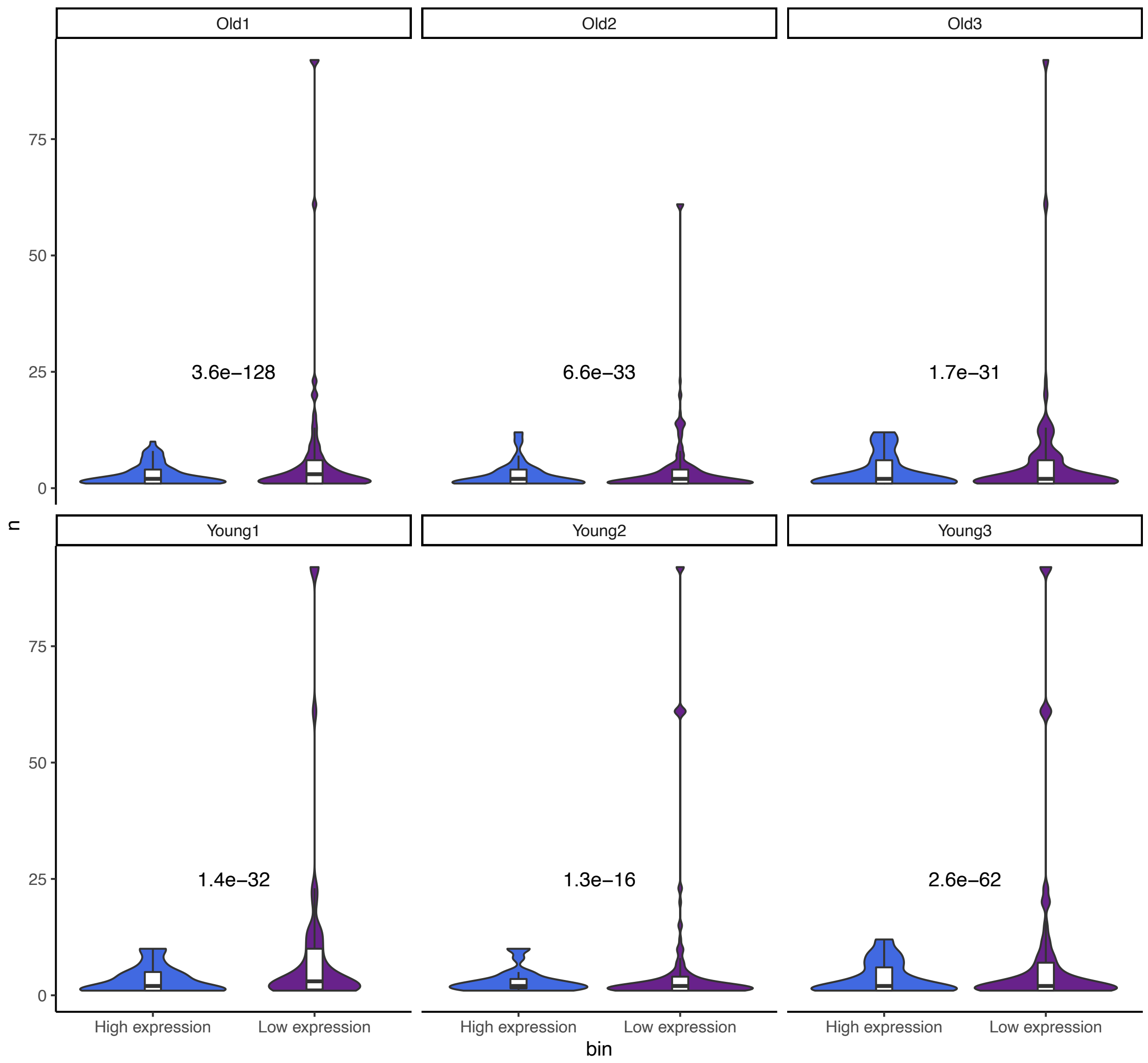


**Supplemental Figure 4: Expression vs. number of SNPs detected.**

We average the expression of every gene across every replicate then compared the number of SNPs detected within genes with < mean expression ("Low expression) with genes with expression greater than the mean expression of all genes ("High expression"). There are significantly more mutations in lowly expressed genes in every replicate.

| Replicate | Number of cells | Average UMIs detected per cell | Single cell number of mapped reads | gDNA number of mapped reads | gDNA coverage |
|---|---|---|---|---|---|

| | | | | | |
|---|---|---|---|---|---|
| **Old1** | 10558 | 10659.74 | 410919834 | 100179710 | 104.553 |
| **Old2** | 14289 | 4241.488 | 140702200 | 41647804 | 43.4658 |
| **Old3** | 4014 | 6692.628 | 142874684 | 53553546 | 55.8913 |
| **Young1** | 4000 | 9958.403 | 138231341 | 102792328 | 107.279 |
| **Young2** | 10000 | 13011.76 | 276598073 | 101604985 | 106.04 |
| **Young3** | 9489 | 13014.51 | 327810578 | 41436733 | 43.2455 |

**Supplemental table 1: summary statistics of each replicate and corresponding gDNA file.**

| Gene | FBID_KEY | P value | $Log_2$ fold change | Bonferroni-adjusted P | Cell type | Enriched in age |
|---|---|---|---|---|---|---|
| **RpII215** | FBgn0003277 | 4.12E-09 | 0.3973355 | 6.33E-05 | gsc, early spermatogonia | Young |
| **RpII18** | FBgn0003275 | 8.75E-71 | 0.61734681 | 1.34E-66 | early spermatids | Young |
| **RpII18** | FBgn0003275 | 3.71E-107 | 0.4150375 | 5.70E-103 | early spermatocytes | Young |
| **RpII18** | FBgn0003275 | 5.65E-29 | 0.82130144 | 8.67E-25 | gsc, early spermatogonia | Young |
| **RpII18** | FBgn0003275 | 2.57E-12 | 0.76464915 | 3.95E-08 | late spermatids | Young |
| **RpII18** | FBgn0003275 | 2.56E-179 | 0.4866866 | 3.94E-175 | late spermatocytes | Young |
| **RpII18** | FBgn0003275 | 3.36E-114 | 0.44202857 | 5.15E-110 | late spermatogonia | Young |
| **RpII15** | FBgn0004855 | 3.79E-33 | 0.80140188 | 5.83E-29 | gsc, early spermatogonia | Young |
| **Rpb8** | FBgn0037121 | 1.68E-41 | 1.0299795 | 2.58E-37 | gsc, early spermatogonia | Young |
| **Rpb8** | FBgn0037121 | 2.79E-140 | 0.49454147 | 4.29E-136 | late spermatogonia | Young |
| **Rpb5** | FBgn0033571 | 3.75E-22 | 0.675871 | 5.76E-18 | gsc, early spermatogonia | Young |
| **Mes4** | FBgn0034726 | 3.21E-08 | 0.27180462 | 0.00049248 | gsc, early spermatogonia | Young |
| **DNApol-iota** | FBgn0037554 | 9.08E-08 | 0.25715784 | 0.00139522 | gsc, early spermatogonia | Young |
| **Tctp** | FBgn0037874 | 1.82E-16 | -0.254537 | 2.80E-12 | late spermatogonia | Old |
| **RpS3** | FBgn0002622 | 1.11E-07 | -0.2738453 | 0.00171142 | early spermatocytes | Old |
| **RpS3** | FBgn0002622 | 1.02E-33 | -0.400618 | 1.56E-29 | late spermatogonia | Old |
| **Rad23** | FBgn0026777 | 6.71E-09 | -0.2740551 | 0.000103 | early spermatocytes | Old |
| **Rad23** | FBgn0026777 | 1.30E-06 | -0.3137124 | 0.02001701 | gsc, early spermatogonia | Old |
| **Pop2** | FBgn0036239 | 3.95E-17 | -0.4439238 | 6.07E-13 | early spermatids | Old |
| **Polr2I** | FBgn0004855 | 4.11E-37 | -0.9381206 | 6.31E-33 | gsc, early spermatogonia | Old |
| **Polr2I** | FBgn0004855 | 4.64E-41 | -0.2622246 | 7.12E-37 | late spermatogonia | Old |
| **Polr2H** | FBgn0037121 | 3.63E-15 | -0.3432375 | 5.58E-11 | early spermatocytes | Old |
| **Polr2H** | FBgn0037121 | 1.04E-46 | -1.27946 | 1.60E-42 | gsc, early spermatogonia | Old |
| **Polr2H** | FBgn0037121 | 5.90E-33 | -0.3419196 | 9.06E-29 | late spermatocytes | Old |
| **Polr2H** | FBgn0037121 | 1.55E-88 | -0.5176787 | 2.37E-84 | late spermatogonia | Old |
| **Polr2F** | FBgn0003275 | 8.72E-30 | -0.5970404 | 1.34E-25 | early spermatocytes | Old |
| **Polr2F** | FBgn0003275 | 5.58E-51 | -1.3372644 | 8.56E-47 | gsc, early spermatogonia | Old |
| **Polr2F** | FBgn0003275 | 9.39E-29 | -1.2519184 | 1.44E-24 | late spermatids | Old |
| **Polr2F** | FBgn0003275 | 8.57E-121 | -0.9244289 | 1.32E-116 | late spermatocytes | Old |

| | | | | | | |
|---|---|---|---|---|---|---|
| **Polr2F** | FBgn0003275 | 1.55E-130 | -0.6554289 | 2.38E-126 | late spermatogonia | Old |
| **Polr2F** | FBgn0003275 | 0 | -1.2930938 | 0 | early spermatids | Old |
| **Polr2E** | FBgn0033571 | 1.67E-34 | -0.9207733 | 2.56E-30 | gsc, early spermatogonia | Old |
| **Polr2A** | FBgn0003277 | 1.46E-19 | -0.6032016 | 2.24E-15 | gsc, early spermatogonia | Old |
| **PolD3** | FBgn0283467 | 1.63E-13 | -0.3674564 | 2.51E-09 | gsc, early spermatogonia | Old |
| **CycG** | FBgn0039858 | 4.03E-46 | -0.4248538 | 6.20E-42 | early spermatids | Old |
| **CG32756** | FBgn0052756 | 3.52E-08 | -0.3531289 | 0.00054089 | gsc, early spermatogonia | Old |
| **Caf1-55** | FBgn0263979 | 2.79E-10 | -0.4207472 | 4.29E-06 | gsc, early spermatogonia | Old |
| **14-3-3epsilon** | FBgn0020238 | 2.41E-07 | -0.2607687 | 0.00370001 | early spermatocytes | Old |
| **14-3-3epsilon** | FBgn0020238 | 2.29E-10 | -0.5121799 | 3.51E-06 | gsc, early spermatogonia | Old |
| **14-3-3epsilon** | FBgn0020238 | 1.76E-27 | -0.3352841 | 2.71E-23 | late spermatogonia | Old |

**Supplemental Table 2: Table of differentially expressed genome maintenance genes.**

These were calculated by splitting the main Seurat object into each cell type and setting the Idents to “age”.

| Gene | P value | $Log_2$ fold change | Bonferroni-adjusted P | Enriched in fly age | Cell type | Gene class |
|---|---|---|---|---|---|---|
| **dmel-testis-AG-merged.15917** | 3.38E-49 | 0.29428559 | 5.33E-45 | Young | Early spermatids | segregating |
| **dmel-testis-AG-mergedplus.2408** | 5.55E-20 | 0.29227805 | 8.75E-16 | Young | Early spermatocytes | segregating |
| **dmel-testis-AG-mergedminus.14760** | 3.92E-09 | 0.3235566 | 6.18E-05 | Young | Late spermatogonia | segregating |
| **dmel-testis-AG-merged.15917** | 1.70E-16 | 0.26208888 | 2.67E-12 | Old | Late spermatids | segregating |
| **dmel-testis-AG-mergedplus.3086** | 6.28E-195 | 1.28564792 | 9.90E-191 | Old | Early spermatids | fixed |
| **dmel-testis-AG-merged.15917** | 1.17E-40 | 0.33935426 | 1.84E-36 | Old | Early spermatids | fixed |
| **CG43760** | 8.56E-87 | 0.5905714 | 1.35E-82 | Young | Early spermatids | fixed |
| **CG43750** | 3.19E-46 | 0.25678987 | 5.02E-42 | Young | Early spermatids | fixed |
| **CG43449** | 3.44E-41 | 0.2949748 | 5.42E-37 | Young | Early spermatids | fixed |
| **CG44174** | 2.47E-149 | 1.19382429 | 3.89E-145 | Old | Early spermatocytes | fixed |
| **CG44227** | 2.61E-27 | 0.29089427 | 4.12E-23 | Old | Early spermatocytes | fixed |
| **CG43760** | 1.16E-10 | 0.26595429 | 1.83E-06 | Young | Early spermatocytes | fixed |
| **CG43750** | 8.31E-154 | 0.93040937 | 1.31E-149 | Old | Cyst cells | fixed |
| **CG43449** | 3.10E-91 | 0.74957663 | 4.89E-87 | Old | Cyst cells | fixed |
| **CG43760** | 5.47E-47 | 0.96654881 | 8.62E-43 | Old | Late spermatocytes | fixed |
| **CG43750** | 1.01E-24 | 0.94625088 | 1.59E-20 | Old | Epithelial cells | fixed |
| **CG43760** | 1.27E-08 | 0.4154728 | 0.00019959 | Old | Epithelial cells | fixed |
| **CG43760** | 5.91E-70 | 1.07260707 | 9.31E-66 | Old | Hub cells | fixed |
| **CG43750** | 1.78E-25 | 0.44217757 | 2.81E-21 | Old | Hub cells | fixed |
| **CG43760** | 1.26E-48 | 0.99877606 | 1.98E-44 | Old | Late spermatogonia | fixed |
| **dmel-testis-AG-merged.2415** | 4.16E-17 | 0.44607375 | 6.55E-13 | Old | Late spermatogonia | fixed |
| **CG43760** | 5.80E-42 | 1.28883889 | 9.15E-38 | Old | Late spermatids | fixed |

| | | | | | | |
|---|---|---|---|---|---|---|
| **CG44329** | 1.08E-15 | 0.29891384 | 1.71E-11 | Old | Late spermatids | fixed |
| **CG43760** | 6.07E-27 | 1.9003241 | 9.56E-23 | Young | Late spermatids | fixed |
| **dmel-testis-AG-merged.7007** | 1.94E-11 | 0.91881875 | 3.06E-07 | Young | GSC, Early spermatogonia | fixed |

**Supplemental Table 3: Differentially expressed *de novo* genes.** Unannotated *de novo* transcripts contain the string "dmel-testis-AG-merged" and correspond to entries in our custom reference gtf file using *de novo* transcripts published in our previous work (Witt et al., 2019).

| Name | P value | $Log_2$ fold change | Bonferroni-adjusted P | Enriched in fly age | Cell type |
|---|---|---|---|---|---|
| **DM412** | 4.45E-17 | 0.28765355 | 7.02E-13 | Old | Early spermatids |
| **Jockey2** | 5.23E-28 | 0.40396657 | 8.25E-24 | Old | Early spermatocytes |
| **Jockey2** | 2.63E-37 | 0.60405864 | 4.15E-33 | Young | Early spermatocytes |
| **BS2** | 4.30E-20 | 1.04723415 | 6.78E-16 | Young | Early spermatocytes |
| **DMRT1B** | 1.61E-79 | 0.50192081 | 2.53E-75 | Old | Cyst cells |
| **DOC** | 1.37E-62 | 0.26769752 | 2.16E-58 | Old | Cyst cells |
| **Jockey2** | 9.17E-62 | 0.26235339 | 1.45E-57 | Old | Cyst cells |
| **TAHRE** | 3.06E-61 | 0.28940335 | 4.82E-57 | Old | Cyst cells |
| **DMRT1B** | 4.01E-53 | 0.2895345 | 6.32E-49 | Old | Cyst cells |
| **HETA** | 2.91E-52 | 0.34062078 | 4.59E-48 | Old | Cyst cells |
| **TRANSIB2** | 2.63E-36 | 0.87271865 | 4.14E-32 | Old | Late spermatocytes |
| **DOC** | 3.39E-33 | 0.7125252 | 5.34E-29 | Old | Late spermatocytes |
| **DMRT1B** | 2.44E-24 | 0.54239771 | 3.84E-20 | Old | Late spermatocytes |
| **Jockey2** | 5.76E-11 | 0.29940896 | 9.07E-07 | Old | Late spermatocytes |
| **DM412** | 1.74E-07 | 0.29024674 | 0.00273574 | Old | Late spermatocytes |
| **TAHRE** | 2.69E-09 | 0.37897876 | 4.24E-05 | Old | Epithelial cells |
| **DOC** | 6.17E-32 | 0.53975512 | 9.73E-28 | Old | Hub cells |
| **Jockey2** | 1.91E-16 | 0.28563862 | 3.00E-12 | Old | Hub cells |
| **Jockey2** | 5.22E-08 | 1.12125594 | 0.00082305 | Young | Late spermatogonia |
| **DOC** | 3.03E-07 | 1.88372443 | 0.00477686 | Young | Late spermatogonia |
| **BS2** | 7.18E-19 | 0.3484718 | 1.13E-14 | Old | Late spermatids |
| **DMRT1B** | 4.46E-16 | 0.39895427 | 7.02E-12 | Old | Late spermatids |
| **DOC** | 1.33E-13 | 0.42069063 | 2.09E-09 | Old | Late spermatids |
| **DM412** | 1.53E-11 | 0.29732294 | 2.41E-07 | Old | Late spermatids |
| **Jockey2** | 3.01E-10 | 0.56404339 | 4.74E-06 | Old | GSC, Early spermatogonia |

**Supplemental Table 4: Table of differentially expressed transposable elements.**
